# Population genomic insights into syntrophic symbioses between marine anaerobic ciliates and intracellular methanogens

**DOI:** 10.1101/2025.07.30.667679

**Authors:** Johana Rotterová, Corinna Breusing, Ivan Čepička, Roxanne A. Beinart

**Affiliations:** Graduate School of Oceanography, University of Rhode Island, Narragansett, Rhode Island, USA; Department of Zoology, Faculty of Science, Charles University, Prague, Czech Republic

**Keywords:** symbioses, host specificity, phylogenomics, mitochondrial genome, WGA

## Abstract

Symbiotic interactions are an ecologically and evolutionary significant phenomenon pertaining to virtually every organism on Earth. For eukaryotes inhabiting extreme environments, syntrophic symbioses with microbes may be key to successfully colonizing new niches, such as globally expanding oxygen-depleted habitats. Multi-domain symbioses between microbial eukaryotes and intracellular methanogenic archaea are crucial to understanding the origins and mechanisms of eukaryotic anaerobiosis. Nearly all anaerobic ciliates, ecologically important protists found in diverse oxygen-depleted environments, host methanogenic endosymbionts, sometimes alongside bacterial partners, that facilitate their anaerobic metabolism. Although vertical symbiont transmission necessarily occurs during ciliate cell division, symbionts might occasionally be acquired horizontally. However, patterns of host-symbiont specificity and intraspecific variability remain poorly understood. Here, we present the first intra-specific genomic analysis of both host and symbionts in such partnerships, providing key insights into the fidelity of eukaryotic-prokaryotic liaisons in anoxia. We assessed the symbiont-host co-diversification and genetic variation across eleven populations of a single undescribed *Metopus* species hosting *Methanocorpusculum* cultured from intertidal sediment locations separated by meters to 1000s of kilometers. Our results show incongruency in host mitochondrial and symbiont phylogenies, indicating a mixed transmission mode. On a genomic level, both host and symbiont populations formed distinct location-specific clusters exhibiting no signs of isolation-by-distance. Instead, ecological factors appear to have driven population genomic divergence at least partly and likely led to differences in metabolic traits. Symbiont comparative and population genomics enable us to further comprehend the complex nature of these multi-partner syntrophic symbioses, crucial to interpreting cell-cell interactions across the domains of life.

## Introduction

It is becoming increasingly clear that microbe-microbe symbioses are ubiquitous and significant across diverse habitats [1–3]. These associations are commonly based on the syntrophic transfer of metabolic substrates between partners (i.e., cross-feeding), which allows the participants to expand into new niches at both ecological and evolutionary timescales [4]. In many cases, these syntrophies contribute to fundamental ecosystem processes by consuming or excreting biogeochemically relevant compounds (e.g., consumption of methane by ANME archaea/sulfate-reducing bacteria symbioses) [5] or are important primary producers (e.g., phytoplankton-diazotroph symbioses) [6]. Many of these associations are obligate and, consequently, must be maintained across generations, either by direct vertical transmission to progeny from parental cells, by horizontal re-acquisition of symbionts from the environment or a combination of both [7]. However, compared to what is known about symbiont inter-generational transmission in multi-cellular hosts like plants and animals, there is a large gap in knowledge regarding these processes in partnerships between microbial eukaryotes (protists), bacteria, and archaea. Both strategies have been recorded in microbe-microbe symbioses (e.g., [8, 9]), and each likely has different but profound impacts on the ecology and evolution of each partner. Given the importance of microbe-microbe symbioses especially in aquatic environments, a better knowledge of partner specificity and symbiont transmission is critical to understand the persistence of these partnerships and how they affect the ecology and evolution of microbial communities. Symbiotic partnerships of unicellular eukaryotes with bacteria and archaea in O_2_-depleted and anoxic habitats are thought to be widespread and represent an adaptive solution to the anaerobic challenges in these harsh environments [10–15]. Compared to numerous well-studied associations between eukaryotes and intracellular bacteria (see [3, 16]), the diversity of archaeal endosymbionts among eukaryotes seems to be uncharted territory, with only two methanogenic archaeal classes confirmed to form intracellular symbioses – Methanobacteria and Methanomicrobia [17–20]. Despite the relative rarity of known archaeal endosymbioses, nearly all anaerobic ciliates studied to date have been observed to host intracellular methanogenic, archaeal endosymbionts, sometimes with additional bacterial partners, that facilitate their anaerobic metabolism [18, 20–22]. Anaerobic ciliates are fast-moving, heterotrophic grazers that rely on fermentation in their mitochondria-related organelles (MROs) to satisfy their energetic needs. The low energy yield of fermentation is overcome through syntrophic partnership with intracellular methanogens that consume ciliate-produced fermentative end-products (i.e., hydrogen) and, thus, make fermentation more thermodynamically favorable while also releasing methane gas [23, 24]. In particular, relationships with methanogenic symbionts of the archaeal genera *Methanobacterium*, *Methanoregula*, and *Methanocorpusculum* appear to be common among anaerobic ciliates [20, 25]. In some marine habitats, protist-associated methanogens can account for a significant portion of methane production, a climate active compound [22, 26, 27]; thus, these ciliate-archaea symbioses have the potential to contribute significantly to the oceanic methane cycle.

Although there are increasing efforts to understand ciliate-archaea symbioses, we still know very little about how the symbionts are transmitted between generations. Past experiments have shown that the symbionts are distributed to both daughter cells during host cellular division and sometimes they divide synchronously with the host, as in the case of *Plagiopyla frontata* [28]. In addition, symbionts were found in the resting cysts of their hosts [29], which all points to vertical transmission, from parent to daughter cells. However, genomic analyses of two methanogenic symbionts indicate very minor genome reduction [21, 30], unlike what is usually seen in obligate bacterial symbionts [31]. This suggests that methanogenic archaea were only recently acquired as intracellular symbionts or counteract ongoing genome reduction through occasional horizontal transmission and recombination. In fact, host switching has been implied in a variety of cases [20, 25, 28, 32, 33]. For example, a recent phylogenetic study by Schrecengost *et al.* [32] showed that the *Methanocorpusculum* genus, an ancestrally endobiotic archaeon frequently found in animal digestive tracts and wastewater [34, 35], is commonly harbored by two unrelated lineages of marine, obligately anaerobic ciliates (Armophorea and Plagiopylea). While the *Methanocorpusculum* symbionts appear to be distinct in most host species, they are not entirely group-specific, suggesting past symbiont swapping and replacements. However, these conclusions are based on ribosomal and intergenic spacer marker gene analyses, which offer limited phylogenetic resolution. Genome-level analyses are necessary for a robust investigation of co-diversification and strain-level variation among *Methanocorpusculum* endosymbionts. Additionally, while previous studies have indicated symbiont swapping and replacement at the species level, signatures of co-diversification consistent with vertical transmission and patterns of symbiont variation have not yet been investigated at the subspecies (population) level.

To address this question, we assessed symbiont-host co-diversification among eleven populations of a single, undescribed species of *Metopus* (Armophorea), previously called *Metopus* sp. 1 [32], and its *Methanocorpusculum* endosymbionts using single-cell genome sequencing. To better understand how genetic and functional diversity is distributed across varying geographic locations and distances, we further explored genomic structure based on single nucleotide polymorphisms and indels and inferred population-level differences in metabolic pathways.

## Materials and Methods

### Sample collection and cell culture

Eleven populations of an undescribed species of a euryhaline anaerobic ciliate, *Metopus* sp. 1 (Armophorea, SAL), previously confirmed to host archaeal endosymbionts from the genus *Methanocorpusculum* [30, 32], were collected from marine and brackish sediments across various geographic scales (Table 1). Ciliate populations were assigned into genus *Metopus* and undefined species rank based on morphology and 18S rRNA gene analysis (Supplemental Methods and Results). This undescribed *Metopus* species clusters within the “marine metopid” clade that also includes the closely related species *Metopus contortus,* as well as *M. parapellitus, M. paraes, M. spiculatus*, and *M. paravestitus* (Supplementary Figures S2 in [32] and S7 in [25]). Specifically, four populations of *Metopus* sp. 1 (FRESH25B, FRESH22, FRESH24, FRESH26) were isolated from the same site and sampling spot, just a meter apart in the outlet of Siders Pond (Falmouth, MA, USA). Three populations (SALT15A, SALT16B, SALT210) were isolated from the same site but different sampling spots ca. 30 meters apart in Salt Pond (Falmouth, MA, USA). Another three populations (BI27A, BLAMEL4B, JUMA2M) were isolated from different sites of the same area within New England (RI, USA), with the BI27A isolation site being ∼30 km apart from the JUMA2M and BLAMEL4B isolation sites (∼5 km apart from each other). All nine populations except JUMA2M were established into a long-term, monociliate culture. The JUMA2M metopid population was isolated from a fresh sample, along with one population (JUMA2P) of an unrelated host ciliate, *Plagiopyla* (Plagiopylea, CONThreeP), confirmed to also host *Methanocorpusculum* [32], for comparison (Table 1). Finally, an eleventh population (FARO3) of *Metopus* sp. 1 from a very geographically distant site in Portugal was selected (Table 1).

Ciliates were isolated and cultivated as described previously [32]. Briefly, 50-ml of intertidal sediments were sampled from marine or brackish environments with fluctuating salinity (Table 1) [36–38]. 1-ml was immediately inoculated into 15-ml Falcon tubes containing 9-ml of Sonneborn’s *Paramecium* medium (varying ratios of ATCC medium 802 and ATCC medium 1525), maintained in a monoeukaryotic polyxenic culture at 23°C, and subcultured by transferring 1-ml of cell inoculum into fresh media at two-week intervals. Each original sediment sample was inoculated into media across a range of salinities, but only successful cultures were retained in the end. Ultimately, our efforts yielded *Metopus* sp. 1 strains growing under predominantly brackish conditions at salinities ranging from about 15 to 30 ppt (Table 1).

### DNA extraction, whole genome amplification and sequencing

DNA from single cells was isolated and amplified using the REPLI-g Advanced DNA Single Cell Kit (Qiagen, Hilden, Germany). Briefly, five to eight individual cells of each studied ciliate population were handpicked from monoeukaryotic cultures, with the exception of JUMA2M and JUMA2P, which were handpicked directly from sediment samples. After removal from the culture or sediment, single cells were each washed three times in sterile media, starved for about two hours to remove any contamination from prey microbes, and washed three times again. Subsequently, each single cell was placed into storage REPLI-g buffer and further processed following manufacturer’s instructions to amplify the whole genome. Amplification of DNA was verified via agarose gel electrophoresis. The DNA concentrations were measured on a Qubit instrument (Agilent, Santa Clara, CA, USA), and the products were purified with ExoSAP-IT (Thermo Fisher Scientific, Waltham, MA, USA). Single-cell libraries were prepared at Psomagen Inc. (Rockville, MD, USA) with the plexWell96 Library Prep Kit (seqWell, Beverly, MA, USA) and sequenced on a NovaSeq 6000 platform (Illumina Inc., San Diego, CA, USA) with a 2×150-bp paired-end protocol to an average depth of 64,267,673 total reads per sample (Table S1).

### Metagenomic binning and generation of symbiont reference pangenome

After quality control, raw reads were trimmed with Trimmomatic [39] and filtered for sequence contaminants through mapping against the human (GRCh38) and PhiX reference genomes. Filtered Illumina reads for each isolated host-symbiont cell were then assembled with metaSPAdes [40], using k-mers from 21 to 121 in 10-step intervals, and binned with MetaBAT2 [41], MaxBin2 [42], and metaWRAP [43]. Binning results were scored with DasTool [44]. The taxonomy of the top-scoring metagenome-assembled genome (MAG) for each holobiont sample was assessed with GTDB-Tk [45] and assembly quality was determined with CheckM [46] based on Methanomicrobiales-specific marker genes. High-quality MAGs with >90% completeness and <5% contamination were selected for further analysis (Table S2). Average nucleotide identities (ANIs) between MAGs were calculated with fastANI to determine species boundaries using a 95% similarity cutoff [47]. All MAGs of the same species were annotated with Prokka [48] and combined into a pangenome with Panaroo [49], using default parameters for gene clustering and a gene re-finding step to locate previously overlooked genes within a radius of 1,000 nt. Spurious annotations were ignored with the --remove-invalid-genes flag. A genus-specific pangenome (including divergent *Methanocorpusculum* species) for phylogenetic analyses was also created by lowering the sequence and family identity thresholds to 0.9 and 0.5, respectively. Genes without functional annotation were further analyzed through BlastP searches [50] against the UniRef90 database with an e-value threshold of 1e-10.

### Host RNA isolation and cDNA sequencing

To generate a host reference transcriptome for population genomic analyses, total RNA from the *Metopus* sp. population FRESH25B was extracted from approximately 0.5-L of a well-grown monoeukaryotic culture using TriReagent (Sigma-Aldrich, St. Louis, MO, USA) and purified using the RNeasy Mini Kit including DNase I treatment (Qiagen, Hilden, Germany). The RNA sample was subsequently sent to Psomagen Inc. (Rockville, MD, USA) for library preparation with the TruSeq Stranded mRNA Kit (Illumina Inc., San Diego, CA, USA). The final RNA-seq library was 150-bp paired-end sequenced on a NovaSeq 6000 platform (Illumina Inc., San Diego, CA, USA) to a depth of about 120 million total reads.

### Assembly of host reference transcriptome

RNA read processing and assembly followed Breusing *et al.* [51]. Briefly, raw reads were quality checked with FastQC (https://github.com/s-andrews/FastQC), trimmed with Trimmomatic [39], and filtered for sequence contaminants, residual rRNA, and bacterial and archaeal sequences. Cleaned reads were error-corrected with Rcorrector [52] and further filtered with TranscriptomeAssemblyTools (https://github.com/harvardinformatics/TranscriptomeAssemblyTools). Assembly was performed with Trinity using the PasaFly algorithm [53]. Assembled contigs were clustered with Cd-hit-EST [54] at a 95% identity threshold to reduce transcript redundancy. Open reading frames (ORFs) were predicted with TransDecoder (https://github.com/TransDecoder/TransDecoder) based on the universal genetic code as suggested for most metopid ciliates [55, 56]. Non-eukaryotic transcripts and transcripts without ORFs were removed from the assembly using a combination of Bowtie2 [57], SAMtools [58], seqtk (https://github.com/lh3/seqtk) and BlobTools [59]. Quality and completeness of the assembly was evaluated with BUSCO based on Alveolata-specific marker genes [60].

### Symbiont and host phylogenomic analysis

To assess host-symbiont co-diversification in the *Metopus*-*Methanocorpusculum* symbiosis we reconstructed phylogenies based on 17 host mitochondrial genes and 1,225 symbiont core genes. Host mitochondrial contigs were obtained through a combination of MITObim [61] and BlastN searches against the sample-specific metagenomes using gene sequences from available ciliate mitogenomes as a query (GenBank: GU057832.1; Dryad doi: 10.5061/dryad.vx0k6djnm - HeterometopusCSS_scaffold_52.gb, Heterometopus_scaffold_11415.gb, Supplementary Data 2 in [62]). Subsequently, iterative BlastN searches in reads were followed by read alignment to the corresponding contig using the “map to reference” option in Geneious Prime (https://www.geneious.com/) to unambiguously extend the longest mitogenome fragment. Initial annotations of mitogenome sequences were obtained using MFannot [63] (https://megasun.bch.umontreal.ca/apps/mfannot/). Predicted genes recovered by MFannot (including hypothetical proteins) were individually checked to confirm their identity and to exclude possible bacterial contamination using Blast searches against the nr/nt database. Final mitogenomes were annotated with GeSeq [64] and the final set of homologous protein-coding, ribosomal and tRNA genes was aligned with MAFFT [65]. A concatenated symbiont core gene alignment was obtained as part of the genus-specific pangenome reconstruction process in Panaroo (see above). A multi-gene phylogeny for the symbiont and individual gene trees for the host were reconstructed in RAxML [66] under the GTR + I + Γ model. The host gene trees were reconciled into a species tree with ASTRAL-Pro [67]. Final maximum-likelihood trees for both host and symbiont were midpoint-rooted with Phangorn [68], plotted with GGtree [69] in R [70], and graphically edited in Inkscape (https://inkscape.org).

### Variant identification in host and symbiont populations

To assess genomic variation among host and symbiont populations, we identified single nucleotide polymorphisms and (if possible) indels across holobiont samples as previously described [51, 71]. As the SALT15A population hosted a different symbiont species and the taxonomic status of the host could not be unambiguously determined, it was removed from all further analyses. Filtered reads from each sample were mapped against the corresponding host transcriptome and symbiont pangenome with Bowtie2 [57] in very sensitive local mode. PCR and optical duplicates were removed with MarkDuplicates (https://github.com/broadinstitute/picard), while LoFreq [72] was applied to realign reads around indel regions and recalibrate base score qualities.

Host population genomic variation was assessed in Angsd [73] by evaluating genotype likelihoods based on an inbred model [74]. We excluded all sites with mapping qualities <30, base qualities <20, and minimum minor allele frequencies <0.01 as well as sites with significant strand or heterozygote bias (p = 0.05), and low probability of being variable. We also discarded improperly paired reads, regions with excessive mismatches and variants close to indel regions. Putative paralogous variants were removed by ignoring reads with non-unique mappings and by setting a threshold on mapping depth based on the average coverage distribution. Genetic distances between samples were obtained by inferring a pairwise covariance matrix.

Symbiont population genomic variation was analyzed with FreeBayes [75] using the --pooled-continuous option. To prevent bias in variant recovery due to uneven read depth between samples, we normalized all samples to the lowest sample-specific coverage (about 15x) prior to analysis. Variant calls were retained if they had a minimum base quality of 20, a minimum mapping quality of 30, a low probability of strand bias (SRP > 5 && SAP > 5 && EPP > 5), no direct proximity to indels (5 bp), and no excessive read depths. In addition, we only included sites and samples that had observations for at least 75% of data points. Consensus haplotype information (as a proxy for the dominant symbiont strain) was obtained with the --extract-FORMAT-info option in VCFtools [76].

The genetic structure of host and symbiont populations was determined through principal-component and coordinate analyses in R, correcting for negative eigenvalues with the method by Cailliez [77].

### Functional variation between symbiont populations

To determine potential functional differences among symbiont populations we assessed KEGG module completeness using the “anvi-estimate-metabolism” program in Anvi’o [78] based on the pathwise completion score. Principal component plots of module completeness scores were generated in R and used for the identification of broader groups for functional enrichment analysis. We defined two multi-population clusters based on the PCA results: 1) FRESH22, FRESH24, FRESH25B, FRESH26, SALT210, BLAMEL4B, 2) BI27A, JUMA2M, and two single-population clusters: 3) FARO3, 4) SALT16B. Enrichment analyses were run for KEGG modules, KOfam annotations and COG20 annotations using the “anvi-compute-metabolic-enrichment” program with default settings.

## Results

### Methanocorpusculum species show host-specific associations and evidence for a mixed transmission mode

Metagenomic binning recovered one high-quality *Methanocorpusculum* symbiont MAG from each ciliate cell, with completeness scores of 90.94–99.42% and maximum contamination levels of 2.41% (Table S2). Symbiont MAGs varied in size from 1.45–2.08 Mb and contained 1,589– 2,446 predicted protein-coding genes, with GC content ranging from 49.33–51.04% (Table S2), as in multiple other host-associated and free-living species of *Methanocorpusculum* [30, 35, 79]. The average nucleotide identity (ANI) between MAGs from different populations ranged from 98.4% to 100%, except in the case of SALT15A and JUMA2P (Table S3). MAGs of these populations showed lower ANIs of only 85.10–89.59% and 85.10–86.76%, respectively, to MAGs from other host populations. ANIs within each population were over 99% for all comparisons. Based on classifications by the Genome Taxonomy Database, most symbionts are conspecific with *Methanocorpusculum* sp003315675, while the SALT15A and JUMA2P populations are more closely related to *Methanocorpusculum labreanum* and *Methanocorpusculum parvum*, respectively (Table S2). These results indicate that the recovered MAGs comprise three *Methanocorpusculum* species that are hosted by two different ciliate genera - one *Methanocorpusculum* is associated with *Plagiopyla* sp. (JUMA2P) and two other closely related species (SALT15A, all others) are associated with *Metopus* sp.

In accordance with these results, phylogenomic analyses based on 1,225 symbiont core genes show that on the mid-point rooted tree, most symbionts form a fully supported single species cluster hosted by *Metopus* sp. 1, while *Methanocorpusculum* associated with SALT15A is a sister clade to *Methanocorpusculum labreanum,* i.e., it is more closely related to a free-living species than to its symbiotic congeners (Fig. 1). Additionally, the *Methanocorpusculum* sp. symbiont isolated from the unrelated *Plagiopyla* host forms an outgroup to all other *Methanocorpusculum* strains included in the analysis. The host mitochondrial phylogeny was incongruent with the symbiont tree (Fig. 1). Decoupling of host and symbiont phylogenies was present both across and within locations, supporting a mixed transmission mode in the *Metopus*-*Methanocorpusculum* symbiosis (Fig. 1).

**Figure 1.**
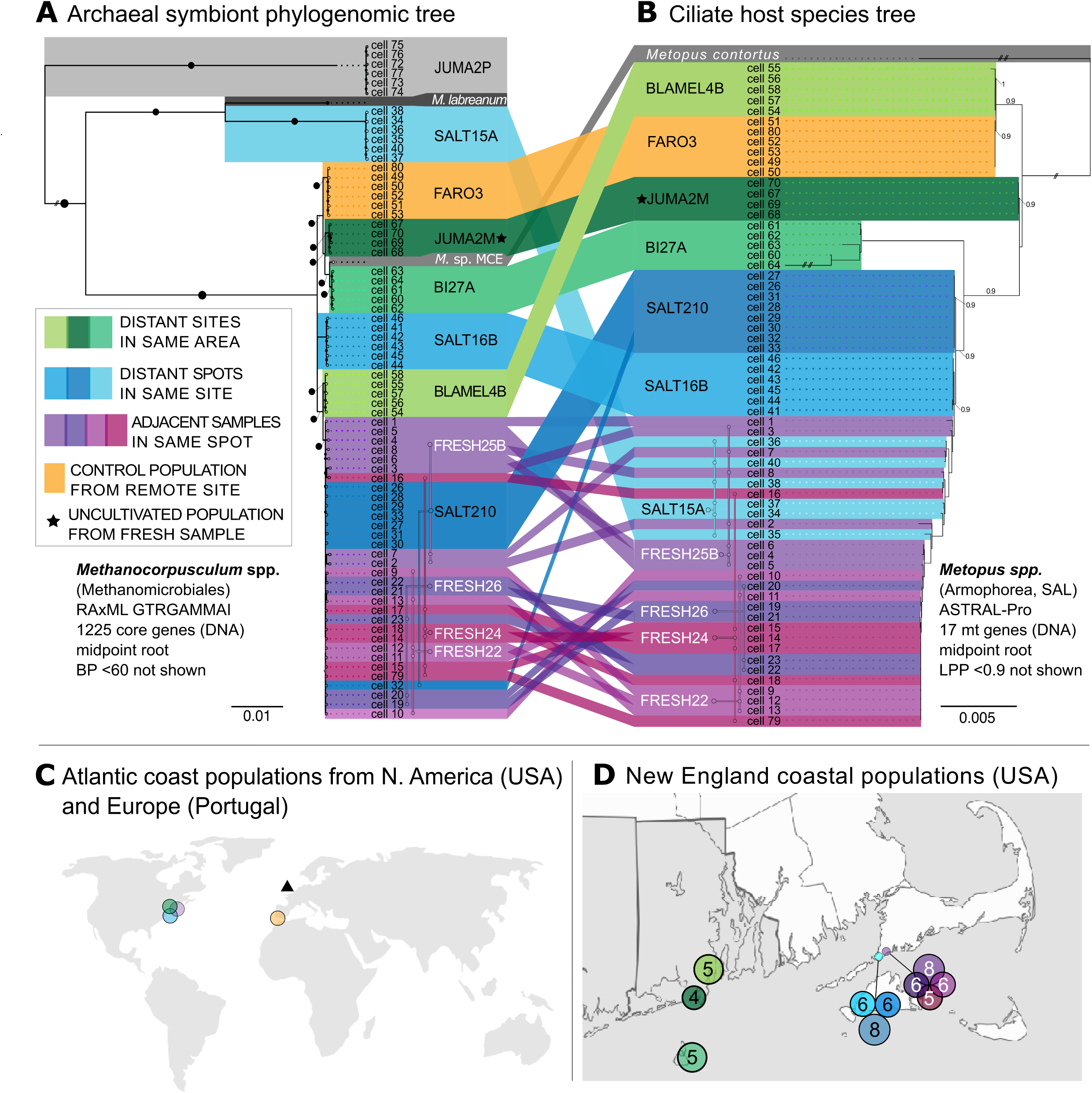
Phylogenomic tree based on 1,225 core gene sequences of the *Methanocorpusculum* symbionts constructed with RAxML (GTRGAMMAI model) (A) and consensus multi-gene mitochondrial phylogeny of host ciliates constructed with ASTRAL-Pro (B). Both trees were mid-point rooted. The values at branches represent statistical support in bootstrap values (BP) or local posterior probabilities (LPP). Symbol legend: large black circle – BP 100; small black circle – BP above 60; // – branch length 50% shortened. A sampling map of ciliate populations is shown for geographic context (C, D), where circles correspond to the herein studied populations, with numbers indicating isolated host cells per population (D). The black triangle represents the isolation site of *Methanocorpusculum* sp. MCE (Lind et al. 2018).

### Host and symbiont populations are structured by locality without evidence for isolation-by-distance

The transcriptome from the FRESH25B population was assembled to serve as reference for the analysis of host population genomic structure. The assembly consisted of 40,151 transcripts (totaling 36.78 Mb) and was approximately 51.50% complete (Table S4). Mapping of host reads from the single cells against the transcriptome reference and filtering of variant sites generated 7,393 SNPs for population genetic analysis. Principal component analyses based on these SNPs indicated that host cells largely clustered by their original sampling locations, which were divided into four broader genetic groups (Fig. 2): 1) FRESH22, FRESH24 (apart from three outliers), FRESH26, and SALT210, 2) FRESH25B, 3) SALT16B, 4) BLAMEL4B, FARO3, JUMA2M, and BI27A. The associations of the FRESH24, SALT210, and SALT16B were inconsistent with the host mitochondrial phylogeny, which implied a closer relationship of FRESH24 and SALT16B with the remaining FRESH samples and a larger phylogenetic distance of SALT210 (Fig. 1). However, both methods clearly implied that genetic or phylogenetic distance was unrelated to geographic distance in *Metopus* ciliates.

**Figure 2.**
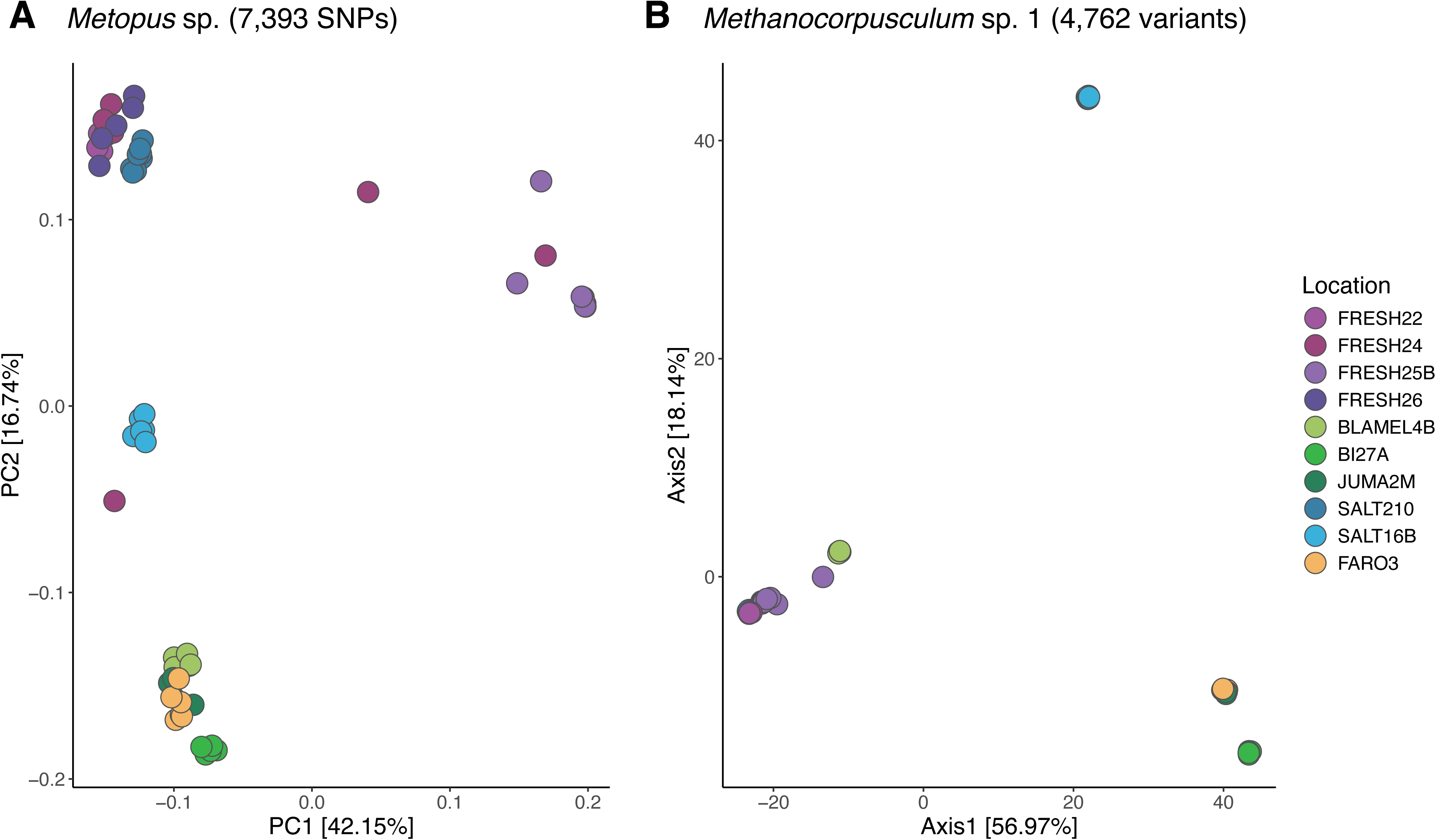
Principal component analysis plot of host (A) and symbiont (B) genetic variants. Dots were slightly jittered to improve visibility of genetic clusters.

For the *Methanocorpusculum* symbiont, we created a species-specific pangenome as reference for population genetic analyses. The pangenome consisted of 1,450 core genes and 2,554 accessory genes, with a total approximate size of 1.60 Mb (Table S4). Mapping against the pangenome and subsequent variant filtering resulted in 4,762 variants (SNPs plus indels) for the assessment of population genomic structure. Like their host counterparts, symbiont samples clustered by locality in principal component analyses (Fig. 2), but the genetic associations among localities, especially for BLAMEL4B and FRESH25B, differed. Symbionts formed three main genetic clusters that were consistent with placements in phylogenetic analyses: 1) all FRESH populations, SALT210 and BLAMEL4B, 2) SALT16B, 3) BI27A, FARO3, and JUMA2M. As in the host, however, the associations among these clusters were independent of geographic distance and clustering patterns did not exactly follow those observed for the ciliate hosts.

### Symbiont populations show divergence in metabolic pathways that is not always related to genetic distance

Ordination analyses of KEGG module completeness indicated the presence of four main metabolic clusters consisting of: 1) all FRESH populations, SALT210, BLAMEL4B, 2) FARO3, 3) SALT16B, and 4) BI27A, JUMA2M (Fig. 3). Although, for the most part, metabolic clustering coincided with genetic clustering based on variant sites, the FARO3 population formed its own metabolic cluster, indicating divergence in the accessory genome. Functional enrichment analyses suggested that over- or under-represented pathways among the four groups primarily concerned the biosynthesis of essential amino acids and vitamins as well as beta-oxidation (Table 2, S5). BI27A and JUMA2M (Group 4) showed significant enrichment in leucine, isoleucine and cobalamin biosynthesis compared to all other groups. Together with FARO3 (Group 2), this group also exhibited over-representation of beta-oxidation related functions. By contrast, histidine biosynthesis and isoleucine biosynthesis from threonine were under-represented in FARO3 and the FRESH/SALT210/BLAMEL4B cluster (Group 1), respectively. Apart from differences in metabolic pathways, *Methanocorpusculum* strains also showed notable contrasts in individual gene functions (Table S5). The most prevalent functional categories among these variable genes were ‘Coenzyme Transport and Metabolism’ and ‘Inorganic Ion Transport and Metabolism.’ For example, together with SALT16B, FARO3 encoded 5-amino-6-(D-ribitylamino)uracil--L-tyrosine 4-hydroxyphenyl transferase, an enzyme involved in the biosynthesis of coenzyme F420 from riboflavin precursors, indicating potential differences between strains in the metabolic subpathways used to generate this cofactor for methanogenesis. In addition, the FRESH/SALT210/BLAMEL4B (Group 1) and BI27A/JUMA2M (Group 4) populations exhibited enrichment in various transporters and a few mobilome related genes that were not found in the other groups. Interestingly, the FARO3 population exclusively contained a peptide methionine sulfoxide reductase (msrA/msrB) involved in oxidative stress response.

**Figure 3.**
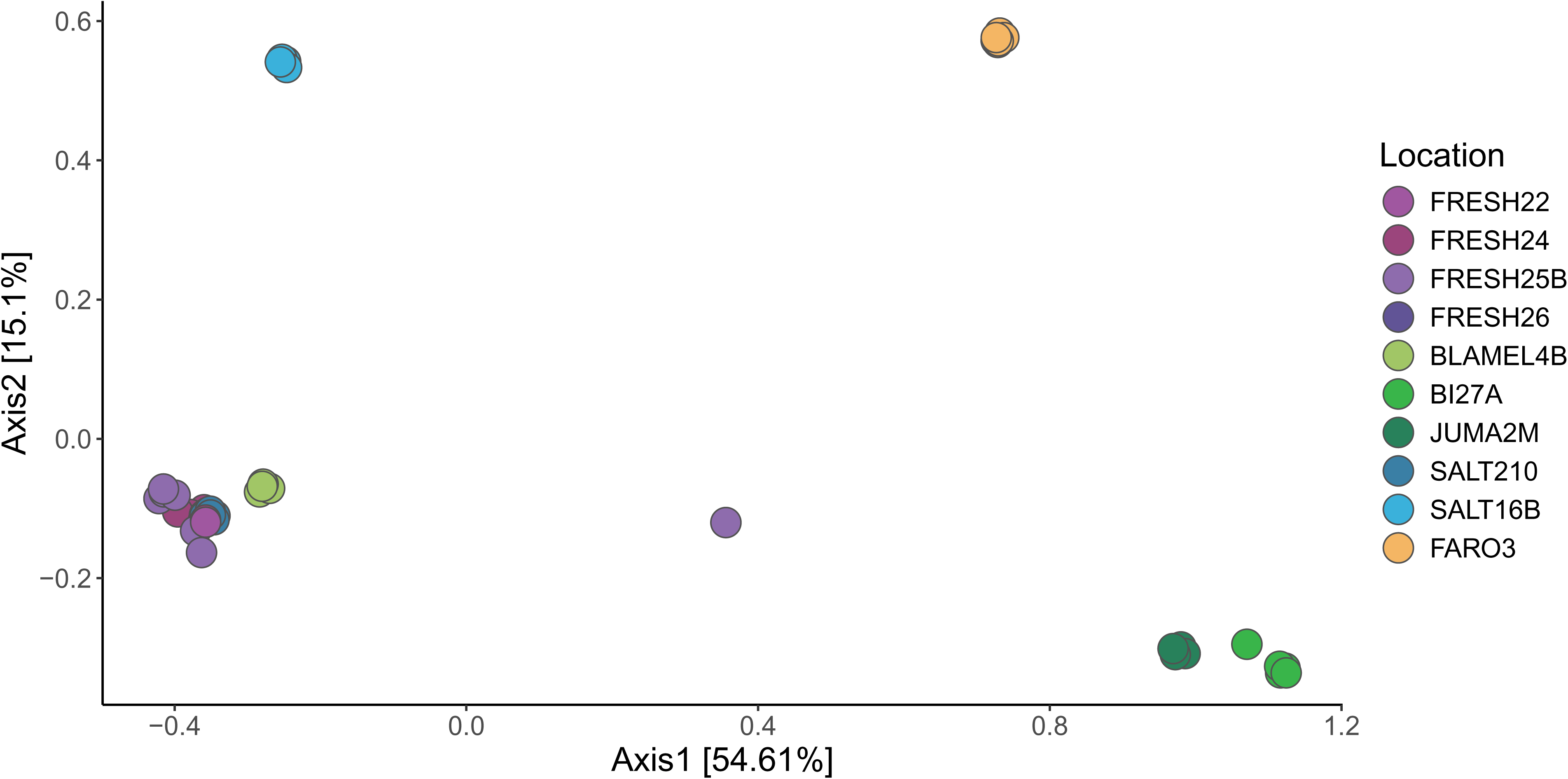
Principal component analysis plot of KEGG module completeness in *Methanocorpusculum* symbiont populations. Dots were slightly jittered to improve visibility of metabolic clusters.

## Discussion

Two unrelated, obligately anaerobic ciliate lineages are known to harbor symbionts of the genus *Methanocorpusculum* [20, 30, 32, 80], a methanogenic archaeal genus otherwise commonly occurring in the gastrointestinal tract of animals and in wastewater environments [34, 35]. Based on small subunit rRNA gene sequences and internal transcribed spacer regions, *Methanocorpusculum* symbionts cluster mostly according to their host species and group taxonomy (Plagiopylea vs. Metopida) [32] and this host fidelity is mostly maintained even when multiple host lineages co-occur in the same location or culture. Above species level, however, incongruent host-symbiont phylogenies have been reported, challenging the assumption that ciliate symbionts are strictly vertically transmitted [32]. Our study expands these conclusions by showing that, below species level, the phylogeny of *Methanocorpusculum* strains does not mirror the MRO phylogeny of their hosts, suggesting that symbiont switches happen on ecological and not just evolutionary timescales. Assuming consistency with the only ciliate (*Paramecium*) in which this has been examined, *Metopus* mitochondria are expected to be passed vertically to daughter cells during binary fission and, unlike ciliate micronuclei, not exchanged during sexual process [81]. While there is no recombination, the selection efficacy in mitochondrial genomes seems to be similar or stronger than in nuclear genomes in all eukaryotes, including ciliates, as shown in the model ciliate *Paramecium* [81]. Thus, if MROs and *Methanocorpusculum* endosymbionts are transmitted with similar fidelity in *Metopus*, we would expect congruence in their phylogenies. Occasional symbiont replacements through horizontal transmission and the resulting opportunities for genetic recombination would explain the observed incongruence in host-symbiont phylogenies and why the genomes of intracellular *Methanocorpusculum* species have remained overall similar in size to their free-living relatives.

Despite lacking the strong genomic and structural reduction typical of intracellular symbionts, the association between the *Methanocorpusculum* symbionts and their hosts seems to be more intimate than in the other two genera of known methanogenic symbionts of anaerobic ciliates. The *Methanocorpusculum* symbiont cells are inserted in stacks with the host’s MRO [30, 32, 82, 83] and are not loosely distributed through the cytoplasm as in other studied ciliate symbioses [20, 21]. The ancestral genome reconstruction of the *Methanocorpusculum* sp. MCE genome showed lower coding densities and much higher rates of pseudogenization compared to its free-living relatives, with ∼5% of genes identified as genetic remnants in the process of being lost [30]. An increased rate of pseudogenization is characteristic for the early stages of adaptation to an intracellular lifestyle [84].

Based on genome-wide variant information, *Methanocorpusculum* symbionts and *Metopus* hosts clustered largely by cultured population. Within host and symbiont populations, genetic differences were mostly insignificant, as cells within populations were more closely related to each other than to cells from other populations, except for the FRESH populations isolated from the same sampling spot in Siders Pond (Falmouth, MA, USA). Intra-population variance within cultivated strains requires careful interpretation and further analyses to exclude the possibility that the very low diversity may be a result of long-term cultivation and, therefore, a population bottleneck. Though ciliates undergo conjugation during their sexual development to increase genetic diversity, this process generally occurs at the end of a phase of exponential vegetative population growth in ciliate cultures when nutrient availability declines [85, 86]. The herein used cultures were re-inoculated before reaching the end of the exponential growth phase, i.e., before cells engaged in conjugation, and, therefore, intra-population diversity might have artificially been reduced. Nevertheless, the uncultivated JUMA2M population, with cells directly picked out of a fresh sediment sample, exhibited similar or even lower intra-population variation compared to the cultivated populations, implying that bottleneck effects and other cultivation-related issues most likely did not significantly bias the results. To fully confirm this conclusion, however, further comparisons among cultivated and freshly isolated samples will be necessary. Moreover, common garden experiments under a variety of environmental conditions will help to understand how ciliate-archaea populations respond to selective pressures in cultivation and how these responses influence measures of genetic differentiation.

The genetic relationships among the culture-specific clusters were not linked to geographic distance, as cultured populations from remote locations were sometimes genetically more similar to each other than populations from the same sampling area. For example, though most *Metopus* hosts from the closely located FRESH populations were very similar, the FRESH25B population seemed less divergent to populations that were 10s to 1000s of kilometers away. Conversely, we observed that some strains from Portugal (FARO3) and the USA (BLAMEL4B, BI27A, and JUMA2M) were very similar despite originating from locations spanning an entire ocean. These findings imply that both ecological factors and long-distance dispersal contribute to the genomic structure of ciliate-archaea symbioses. Although the dispersal mechanisms of free-living ciliates are understudied, these organisms are known to be dispersed by various ways, such as waterbirds and possibly within stress-resistant cysts [87]. There are indications that biogeographic processes influencing microbial groups in island systems depend mostly on efficient dispersal, and are facilitated by a high stress tolerance, leading to easier establishment in novel habitats [88]. Most populations of *Metopus* sp. 1 seem to be euryhaline and can thrive in a wide range of salinities [20, 30, 89], which likely allows these protists to colonize a variety of different environments. Indeed, most of the herein studied populations were isolated from marine or brackish sites with highly variable salinity, such as coastal salt ponds, tidal lagoons, and salt marshes. Unfortunately, we do not have enough information about the environmental conditions of the sampling locations to assess whether and how the population genetic structure observed here reflects adaptation to local habitat characteristics like salinity, nutrient levels, or oxygen concentrations. Given that most cultures were maintained under the same media conditions it is unlikely that the observed patterns of inter-population differentiation were driven by adaptation to long-term cultivation, although future analyses will need to be performed to verify this assumption.

Interestingly, the archaeal symbiont did not show the exact same population structure as the ciliate host. Despite divergence in the host, the *Methanocorpusculum* symbionts of the FRESH populations from Siders Pond were all quite similar, suggesting local symbiont exchange and horizontal transmission at this scale. These findings indicate that ciliate hosts and symbionts are subject to partly differing dispersal processes and/or environmental pressures, which is consistent with a mixed transmission mode where symbionts evolve semi-independently from their hosts [90, 91]. Metabolic differences among symbiont populations mainly concerned the potential for biosynthesis of some essential amino acids and cobalamin, as well as beta-oxidation. Only the JUMA2M and BI27A appeared to be able to synthesize cobalamin, a critical vitamin in the methanogenesis pathway, while all other populations seemed to lack this capacity. Multiple *Methanocorpusculum* populations investigated here were also unable to produce isoleucine, leucine and/or histidine. The lack of functional pathways for the biosynthesis of cobalamin and essential amino acids has previously been reported in the related *Methanocorpusculum* sp. MCE symbiont [30, 92], where critical genes involved in these metabolisms appeared to be pseudogenized [30]. These observations could suggest that *Methanocorpusculum* symbionts have reached different stages of reductive genome evolution, possibly due to differential rates of supplemental horizontal transmission. However, the fact that genes for critical biosynthesis pathways, like chorismate and aromatic amino acids, are also missing in several free-living methanogens (*Methanocorpusculum bavaricum*, *Methanobrevibacter oralis*) implies that the loss of these functions may be common in methanogenic archaea independent of their lifestyle [30]. Given that *Methanocorpusculum* species often occur in nutrient-rich habitats where all necessary vitamins and amino acids are easily accessible, it is likely that reduced selective pressures for gene retention led to loss of these biosynthetic functions in some populations, while others might have evolved alternative metabolic pathways, such as cobalamin-independent systems for methanogenesis [93]. Despite the lack of environmental data for the sampled locations, our gene enrichment analyses point to potential differences in habitat characteristics that might have led to divergence in gene content among *Methanocorpusculum* populations. For instance, we observed variation in genes related to oxidative stress, ion transport, F420 biosynthesis and mobile elements. This could indicate that the sampled populations experience differences in oxygen penetration, nutrient profiles and/or viral communities, a hypothesis that should be investigated through co-registered sampling of holobiont populations and environmental parameters in subsequent studies.

Overall, our results provide new insights into the transmission modes and evolutionary dynamics of aquatic ciliate-archaea symbioses, while raising interesting questions for future research. For example, under which conditions are horizontal transmission and symbiont switching induced? Are there constraints on the type of symbiont replacements and how does this impact genome evolution in the symbiont? Which ecological factors determine the genomic and functional divergence of hosts and symbionts? Answers to these questions will not only advance our understanding of ciliate-archaea symbioses but also shed light on broader evolutionary processes, such as the evolution of the eukaryotic cell.

## Data Availability

Raw metagenomic reads and *Methanocorpusculum* MAGs have been submitted to GenBank under BioProject PRJNA1289166, while RNA-seq reads and the *Metopus* sp. 1 transcriptome assembly have been deposited in PRJNA1289170. New 18S rRNA sequences are available on GenBank under accession numbers PV928686–PV928689.

## Acknowledgments

The authors would like to thank Anna Schrecengost for sampling and cultivation efforts.

## Author contributions

Johana Rotterová: Data collection, Formal analysis, Investigation, Writing – original draft, Writing – review & editing. Corinna Breusing: Formal analysis, Writing – original draft, Writing – review & editing. Ivan Čepička: Resources, Funding acquisition, Writing – review & editing. Roxanne Beinart: Conceptualization, Methodology, Resources, Funding acquisition, Writing – original draft, Writing – review & editing.

