## Supplemental Methods for "Population genomic insights into syntrophic symbioses between marine anaerobic ciliates and intracellular methanogens"

**Supplemental Methods and Results**

### *Organism identification*

Ciliate populations were assigned into genus *Metopus* and undetermined species rank based on morphology and 18S rRNA gene analysis. Specifically, the morphological taxonomy designation is based on the grossly uniform morphology resembling *Metopus contortus* (Quennerstedt, 1867) Kahl, 1932 but fine differences in size and cortical structures observed on live and protargol stained specimens. The 18S rRNA gene sequences of ciliate populations BI27A, BLAMEL4B, FARO3, FRESH24, FRESH26, and FRESH25B were generated during a previous study [1], and sequences from the remaining herein studied populations (FRESH22, JUMA2M, SALT210, SALT16B, and SALT15A) were generated using the same methods of DNA extraction, amplification, and sequencing.

To estimate divergence of the 18S rRNA gene sequences, we calculated pairwise distances with Clustal Omega in Geneious Prime (https://www.geneious.com) [2], and genetic distances (uncorrected p distances) in MEGA11 based on site rate variation modeled with a gamma distribution [3, 4]. The pairwise nucleotide sequence identity ranged from 99% to 100% and genetic distances (uncorrected p distance) were below 0.002 between all populations except SALT15A. The pairwise nucleotide sequence identity between population SALT15A and all others was 97.9-98.6% with p-value 0.004. Using a conservative approach, the SALT15A population was not determined to be a different species, since the sequence divergence observed here is lower than intraspecific divergence of the closely related *Metopus contortus* ciliate (93-98.5% based on 18S rRNA gene sequences available in GenBank).
